# Ark-ADMET: endpoint-specific selection of molecular representations and hyperparameters for gradient-boosted ADMET models under the Therapeutics Data Commons benchmark protocol

**DOI:** 10.64898/2026.10.28.762420

**Authors:** Yuzhu Shi, Haoxi Luo, Feng Liu, Simiao Zhao

## Abstract

Typical submissions to the Therapeutics Data Commons (TDC) absorption, distribution, metabolism, excretion, and toxicity (ADMET) benchmark leaderboard apply a single architecture and a single feature representation across all 22 endpoints, despite substantial heterogeneity in evaluation metric, training-set size, and class balance. Rather than proposing a new architecture, we show that endpoint-specific selection of molecular representations and CatBoost hyperparameters, carried out under the official TDC ADMET benchmark protocol, is a competitive alternative to that uniform recipe. For each endpoint, a fixed CatBoost backbone was paired with configurations chosen from a common candidate set comprising published fingerprints and physicochemical descriptors, truncated three-dimensional PaDEL descriptors, frozen graph isomorphism network (GIN) and MiniMol embeddings, and a convex combination with a shallow network trained on the MiniMol embedding alone. All feature and hyperparameter choices were made on an internal scaffold validation split under a prespecified strict-improvement rule. Against a frozen snapshot of the public leaderboard, the resulting Ark-ADMET family of 22 endpoint-specific models matched or exceeded the best previously listed score on 7 of 21 rankable endpoints and strictly exceeded it on 3. All selected configurations retain the baseline fingerprint-and-descriptor block; optional components differ across endpoints. Taken together, these results show that competitive performance on this benchmark can be achieved without a new architecture or a single optional representation shared across all endpoints.

## 1. Introduction

Drug discovery requires absorption, distribution, metabolism, excretion, and toxicity (ADMET) properties to be assessed across many endpoints. Underlying assays differ in measurement type, dynamic range, class balance, and sample size, so a molecular representation useful for one endpoint is not necessarily useful for another. Public ADMET leaderboard submissions nevertheless commonly apply one architecture and one feature set to every endpoint.

The Therapeutics Data Commons (TDC) ADMET benchmark group provides the basis for comparison.^1^ For each of 22 endpoints it fixes a scaffold split between a combined training–validation table and a held-out test table, ^2,3^ and requires reporting the mean and dispersion across five repeats. Two prior approaches motivate the present work. One prior study introduced the MapLight feature block (concatenated Morgan, Avalon, and ErG fingerprints plus 200 physicochemical descriptors) for use with CatBoost^4,5^ and showed that this model is competitive with, and often better than, deep neural models trained for the same task; appending a frozen graph-neural-network embedding improved performance further.^5^ A subsequent study showed that a compact pretrained graph encoder, kept frozen and followed by a shallow head, transfers across the same 22 tasks.^6^ Both approaches apply one configuration to every endpoint. The present candidate set is therefore anchored on the MapLight feature block and on frozen embeddings of this kind, and additionally includes truncated three-dimensional PaDEL descriptors^7^—which may complement two-dimensional fingerprints for some endpoints—and a convex combination with a shallow network trained on the MiniMol embedding alone, so that the published encoder can be used either as tree features, as a separate predictor, or both.

The contribution of this work is not a new architecture. It is to demonstrate that endpoint-specific selection among these published components, when performed under an internal-validation selection protocol, is a competitive alternative to applying one configuration to every endpoint. Concretely, we ask (i) which of the available representations to keep for each endpoint and (ii) which CatBoost hyperparameters should accompany the representations that are kept. Both questions are selection problems. Selection should not rely on the held-out test set: with 22 endpoints and several binary choices per endpoint, revising decisions after seeing test scores can yield a favorable combination by chance, a recognized risk in model selection.^8^

Candidates were therefore compared on an internal scaffold validation split taken only from the training partition of one random seed. A candidate replaced the current configuration only if it was strictly better on that split; ties kept the current configuration. CatBoost was fitted only on the official training partition and was never fitted to, or stopped on, the official validation partition. After a decision was fixed, models were refitted on each of the five official training partitions, and the official test set was scored once per seed.

Under this selection protocol we obtained Ark-ADMET, a family of 22 endpoint-specific CatBoost models that share the MapLight feature block but differ in three-dimensional descriptors, frozen embeddings, and combination weights. On a frozen leaderboard snapshot, this family matched or exceeded the best previously listed score on 7 of 21 rankable endpoints, showing that a uniform feature recipe is not required for competitive standing on this benchmark. A Chemprop model^9^ was trained separately on the same five official splits and is reported as a separate submission; its predictions are neither used as features nor averaged with those of the trees, and fixing CatBoost for the main analyses is not a claim that tree ensembles outperform message-passing networks. To document the selection process, we also report internally selected changes that decreased held-out performance.

## 2. Methods

### 2.1. Benchmark endpoints and evaluation metrics

The 22 endpoints of the TDC ADMET benchmark group ^1^ are identified below by their TDC dataset names: `Caco2_Wang`, `Bioavailability_Ma`, `Lipophilicity_AstraZeneca`, `Solubility_AqSolDB`, `HIA_Hou`, `Pgp_Broccatelli`, `BBB_Martins`, `PPBR_AZ`, `VDss_Lombardo`, `CYP2C9_Veith`, `CYP2D6_Veith`, `CYP3A4_Veith`, `CYP2C9_Substrate_CarbonMangels`, `CYP2D6_Substrate_CarbonMangels`, `CYP3A4_Substrate_CarbonMangels`, `Half_Life_Obach`, `Clearance_Hepatocyte_AZ`, `Clearance_Microsome_AZ`, `LD50_Zhu`, `hERG`, `AMES`, and `DILI`.

Evaluation metrics follow the benchmark specification. `Caco2_Wang`, `Lipophilicity_AstraZeneca`, `Solubility_AqSolDB`, `PPBR_AZ`, and `LD50_Zhu` are evaluated by mean absolute error (MAE), for which lower values indicate better performance. `CYP2C9_Veith`, `CYP2D6_Veith`, `CYP3A4_Veith`, `CYP2C9_Substrate_CarbonMangels`, and `CYP2D6_Substrate_CarbonMangels` are evaluated by the area under the precision–recall curve (AUPRC). `CYP3A4_Substrate_CarbonMangels` and the remaining classification endpoints are evaluated by the area under the receiver operating characteristic curve (ROC-AUC). `VDss_Lombardo`, `Half_Life_Obach`, `Clearance_Hepatocyte_AZ`, and `Clearance_Microsome_AZ` are evaluated by the Spearman rank correlation coefficient. The metric assigned to each endpoint is also listed in Table 1.

**Table 1.** Held-out test performance on the TDC ADMET benchmark group. Leaderboard values and ranks follow Section 2.8: per-seed rounding to three decimal places, then mean and population standard deviation for leaderboard entries; five-seed values in the table are the unrounded mean and sample standard deviation; ranks are assigned against a frozen leaderboard snapshot. Arrows indicate the direction of improvement. Rank is given as rank / number of entries on the board including this study; “(tied)” indicates agreement to three decimal places with at least one previously listed score. `Bioavailability_Ma` is not ranked (Section 2.8).

| Endpoint | Metric | Leaderboard score (mean ± SD) | Five seeds, unrounded (mean ± SD) | Rank | Best previously listed |
|---|---|---|---|---|---|
| Solubility_AqSolDB | MAE ↓ | 0.741 ± 0.008 | 0.7413 ± 0.0093 | 1 / 19 (tied) | 0.741 |
| HIA_Hou | ROC-AUC ↑ | 0.993 ± 0.002 | 0.9934 ± 0.0018 | 1 / 21 (tied) | 0.993 |
| Clearance_Microsome_AZ | Spearman ↑ | 0.634 ± 0.006 | 0.6345 ± 0.0070 | 1 / 21 | 0.630 |
| PPBR_AZ | MAE ↓ | 7.349 ± 0.025 | 7.3493 ± 0.0281 | 1 / 21 | 7.440 |
| AMES | ROC-AUC ↑ | 0.871 ± 0.001 | 0.8713 ± 0.0015 | 1 / 21 (tied) | 0.871 |
| CYP2C9_Veith | AUPRC ↑ | 0.859 ± 0.003 | 0.8592 ± 0.0035 | 1 / 21 (tied) | 0.859 |
| CYP3A4_Veith | AUPRC ↑ | 0.917 ± 0.002 | 0.9172 ± 0.0019 | 1 / 20 | 0.916 |
| LD50_Zhu | MAE ↓ | 0.570 ± 0.013 | 0.5698 ± 0.0148 | 2 / 23 | 0.552 |
| CYP2D6_Veith | AUPRC ↑ | 0.788 ± 0.004 | 0.7881 ± 0.0044 | 2 / 20 | 0.790 |
| Pgp_Broccatelli | ROC-AUC ↑ | 0.930 ± 0.008 | 0.9299 ± 0.0094 | 3 / 17 (tied) | 0.938 |
| BBB_Martins | ROC-AUC ↑ | 0.917 ± 0.006 | 0.9168 ± 0.0068 | 3 / 27 | 0.924 |
| VDss_Lombardo | Spearman ↑ | 0.681 ± 0.017 | 0.6808 ± 0.0192 | 3 / 20 | 0.713 |
| Half_Life_Obach | Spearman ↑ | 0.548 ± 0.042 | 0.5481 ± 0.0473 | 4 / 21 | 0.576 |
| Clearance_Hepatocyte_AZ | Spearman ↑ | 0.448 ± 0.015 | 0.4479 ± 0.0164 | 4 / 19 | 0.536 |
| Lipophilicity_AstraZeneca | MAE ↓ | 0.479 ± 0.007 | 0.4790 ± 0.0079 | 4 / 22 (tied) | 0.456 |
| hERG | ROC-AUC ↑ | 0.864 ± 0.008 | 0.8641 ± 0.0093 | 5 / 21 | 0.880 |
| CYP3A4_Substrate_CarbonMangels | ROC-AUC ↑ | 0.641 ± 0.023 | 0.6413 ± 0.0261 | 7 / 20 | 0.667 |
| CYP2D6_Substrate_CarbonMangels | AUPRC ↑ | 0.692 ± 0.013 | 0.6922 ± 0.0150 | 8 / 19 | 0.736 |
| DILI | ROC-AUC ↑ | 0.891 ± 0.016 | 0.8907 ± 0.0175 | 9 / 21 | 0.956 |
| Caco2_Wang | MAE ↓ | 0.301 ± 0.009 | 0.3008 ± 0.0099 | 9 / 25 | 0.256 |
| CYP2C9_Substrate_CarbonMangels | AUPRC ↑ | 0.385 ± 0.018 | 0.3846 ± 0.0201 | 11 / 21 | 0.474 |
| Bioavailability_Ma | ROC-AUC ↑ | 0.744 ± 0.026 | 0.7445 ± 0.0290 | not assigned | — |

Chemprop, trained and submitted separately: Lipophilicity\_AstraZeneca MAE 0.412, rank 1 / 22.

### 2.2. Data partitioning and selection protocol

Assignment of molecules to the training–validation table and to the held-out test table is fixed by the benchmark and was not modified. Within the training–validation table, `get_train_valid_split(split_type=“default”)` was called for each of the five random seeds; for this benchmark group, the function performs a scaffold split with partition fractions 0.875, 0.125, and 0. CatBoost was fitted exclusively to the 87.5% training partition of the corresponding seed. For CatBoost, the 12.5% official validation partition was not used for fitting, early stopping, or selection; it was used only for Chemprop checkpoint selection (Section 2.7).

All selection decisions for a given endpoint were made on one internal validation split, constructed by Murcko scaffold partitioning ^2^ of the training partition of seed 1 only; 80% of those molecules were used to fit the candidate configuration and 20% to evaluate it. This internal split is distinct from the official validation partition. A candidate configuration replaced the currently selected one only if it achieved a strict improvement in the endpoint-specific metric on the evaluation portion; ties were resolved in favor of the currently selected configuration. Once a decision was fixed, models were refitted on the training partition of each of the five seeds, and the held-out test set was predicted once per seed (Figure 1A). Seeds 2–5 followed the same 87.5%/12.5% partitioning and were used only for these refits and test evaluations; they did not enter any selection decision.

**Figure 1.**
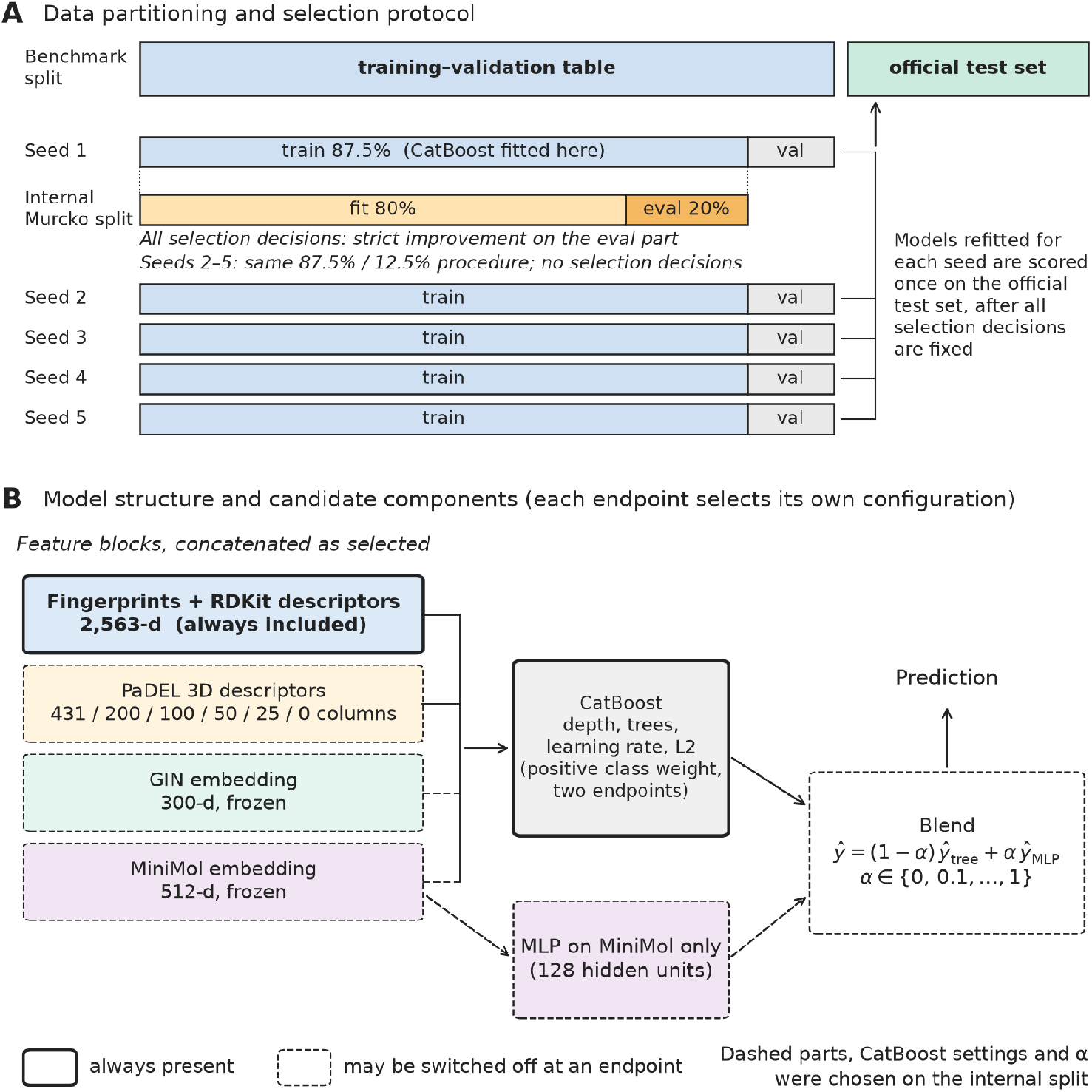
Data partitioning and candidate components. (A) Data partitioning and selection protocol. The scaffold split separating the training–validation table from the held-out test table is fixed by the benchmark. Each of five seeds partitions the training–validation table into training and validation portions at fractions of 0.875 and 0.125, and CatBoost is fitted to the training portion only. All selection decisions are made on an internal Murcko scaffold validation split derived from the training portion of seed 1 (80% fit, 20% evaluation), under a strict-improvement criterion; the official validation portions (grey) play no role in these decisions. The held-out test set is evaluated once per seed after each decision has been fixed. (B) Model structure and candidate components from which each endpoint selected independently. Solid outlines mark components present at every endpoint (the MapLight feature block and CatBoost); dashed outlines mark components that could be switched off. Truncation of the three-dimensional descriptors, concatenation of the GIN and MiniMol embeddings, the CatBoost hyperparameters, the combination weight α, and the positive class weight (evaluated at two endpoints only) were all chosen on the internal validation split.

### 2.3. Candidate molecular representations

All endpoints retain in full the 2,563-dimensional MapLight feature block described by Notwell and Wood: ^5^ hashed Morgan count fingerprints of radius 2 folded into 1,024 bins, ^10^ Avalon count fingerprints folded into 1,024 bins, ^11^ extended reduced graph (ErG) pharmacophore descriptors in 315 bins,^12^ and 200 RDKit physicochemical descriptors.^13^

Three-dimensional descriptors comprise 431 PaDEL-Descriptor columns^7^ computed from a single conformer per molecule. Conformers were generated with ETKDGv3 using a fixed random seed of 1 and relaxed with the Merck molecular force field (MMFF) for at most 200 iterations, with random starting coordinates used as a fallback when embedding failed. Candidate truncations retained the first 431, 200, 100, 50, 25, or 0 columns after ordering by CatBoost feature importance computed on the internal training molecules.

Two pretrained encoders were available as candidates, neither of which was fine-tuned on benchmark labels: the 300-dimensional `gin_supervised_masking` embedding^14^ and the publicly released 512-dimensional MiniMol v1 encoder.^6^ Each embedding was concatenated to the feature vector only for endpoints at which the internal validation split selected it, without additional rescaling (Figure 1B).

### 2.4. CatBoost training

Tree models were CatBoost classifiers and regressors^4^ with `random_strength` set to 2, log loss for classification and MAE for regression. Models were trained for the prescribed number of trees without early stopping. The hyperparameter grid comprised 54 combinations: depth 4, 6, or 8; 500, 1,000, or 2,000 trees; learning rate 0.03 or 0.1; and L2 leaf regularization 1, 3, or 10. The initial grid search evaluated first the configuration reported by Notwell and Wood^5^ (depth 6, 1,000 trees, learning rate 0.03, L2 leaf regularization 3), and ties were resolved in its favor; subsequent searches evaluated the currently selected configuration first, with ties resolved in the same way.

Regression targets were transformed prior to fitting in three steps: subtraction of min(min(*y*), 0) to enforce non-negativity, an optional base-10 logarithmic transformation of the form log_10_(*y* + 1), and standardization to zero mean and unit variance using statistics computed on the training molecules of the corresponding seed. Predictions were transformed back to the original scale by reversing these steps. The logarithmic transformation was applied to `VDss_Lombardo`, `Half_Life_Obach`, `Clearance_Hepatocyte_AZ`, and `Clearance_Microsome_AZ`.

Positive class weighting (`scale_pos_weight`) was evaluated for the two substrate endpoints assessed by AUPRC, both of which exhibited class imbalance, using candidate values of 1, 2, 3, and 5, with the value 1 evaluated first and retained in the event of a tie. A weight of 3 was selected for `CYP2D6_Substrate_CarbonMangels`; `CYP2C9_Substrate_CarbonMangels` retained a weight of 1. All other endpoints used a weight of 1.

### 2.5. MiniMol auxiliary network

Independently of embedding concatenation, a multilayer perceptron (MLP) was trained on the 512-dimensional MiniMol embedding alone. Features were standardized using statistics computed on the training molecules of the corresponding seed. The network had a single hidden layer of 128 units with rectified linear activation and was trained with the Adam optimizer (L2 penalty 10^−4^, initial learning rate 10^−3^) for a maximum of 500 iterations, without early stopping. Regression targets were transformed as described in Section 2.4.

Predictions were formed as the convex combination

*ŷ* = (1 − α) *ŷ*_tree_ + α *ŷ*_MLP_, α ∈ {0, 0.1, …, 1.0}, with α selected on the internal validation split and α = 0 retained in the event of a tie. Molecules that MiniMol could not featurize were scored by the tree alone; this applied to two molecules in the `Solubility_AqSolDB` test set.

### 2.6. Selection sequence

Selection proceeded as an ordered sequence of stages, each applied to the configuration retained by the preceding stage. An initial model was fitted with the published hyperparameters on the MapLight feature block together with all 431 three-dimensional columns, and the 54-combination grid was then searched on that same design matrix. With the selected hyperparameters held fixed, each endpoint was next evaluated under three feature configurations: the MapLight feature block with all three-dimensional columns, the same block with the GIN embedding appended, and the MapLight feature block alone. Endpoints that retained three-dimensional columns subsequently selected a truncation length. MiniMol concatenation was evaluated next, and values of α were then selected on the resulting configuration.

The design matrix changed after the initial grid search for some endpoints. The 54-combination grid was therefore repeated for six of those endpoints: `CYP2C9_Substrate_CarbonMangels`, `CYP2D6_Substrate_CarbonMangels`, `hERG`, `Bioavailability_Ma` (unranked; Section 2.8), `VDss_Lombardo`, and `LD50_Zhu`. `VDss_Lombardo` and `LD50_Zhu` retained their existing hyperparameters. Where hyperparameters changed and the previously selected α was non-zero, α was selected again. For nine other endpoints whose design matrices had also changed, Table 2 reports hyperparameters selected on the initial design matrix (the MapLight feature block plus all 431 three-dimensional columns), marked “earlier.” Seven endpoints underwent no change of design matrix and were not re-searched, since the grid would have been identical. Positive class weighting for the two substrate endpoints assessed by AUPRC was evaluated after this hyperparameter reselection, as described in Section 2.4, and a weight of 3 was adopted for `CYP2D6_Substrate_CarbonMangels`.

**Table 2.** Configurations selected for each endpoint. Hyperparameters are given as depth, number of trees, learning rate, and L2 leaf regularization. “Earlier” indicates that hyperparameters were selected on the MapLight feature block with all 431 three-dimensional columns, prior to the availability of the GIN and MiniMol embeddings; “final” indicates selection on the design matrix used at evaluation. *n* denotes the number of molecules in the training–validation table. α is the combination weight defined in Section 2.5. “Log transform” indicates the base-10 logarithmic target transformation described in Section 2.4.

| Endpoint | n | Hyperparameters | 3D descriptor columns | GIN concatenated | MiniMol concatenated | α | Positive class weight | Log transform | Design matrix |
|---|---|---|---|---|---|---|---|---|---|
| Solubility_AqSolDB | 7,985 | 6, 1000, 0.1, 10 | 50 | no | no | 0.2 | 1 | no | earlier |
| HIA_Hou | 461 | 4, 500, 0.03, 3 | 0 | yes | no | 0.0 | 1 | no | earlier |
| Clearance_Microsome_AZ | 881 | 8, 2000, 0.03, 10 | 431 | yes | yes | 0.0 | 1 | yes | earlier |
| PPBR_AZ | 2,231 | 6, 500, 0.03, 10 | 25 | yes | no | 0.1 | 1 | no | earlier |
| AMES | 5,821 | 8, 2000, 0.03, 10 | 0 | no | yes | 0.2 | 1 | no | earlier |
| CYP2C9_Veith | 9,673 | 6, 1000, 0.1, 10 | 0 | yes | no | 0.2 | 1 | no | earlier |
| CYP3A4_Veith | 9,861 | 6, 2000, 0.03, 10 | 200 | yes | no | 0.1 | 1 | no | earlier |
| LD50_Zhu | 5,907 | 8, 2000, 0.1, 10 | 0 | no | no | 0.2 | 1 | no | final |
| CYP2D6_Veith | 10,504 | 8, 2000, 0.03, 10 | 431 | yes | yes | 0.0 | 1 | no | earlier |
| Pgp_Broccatelli | 973 | 8, 500, 0.03, 10 | 0 | no | no | 0.0 | 1 | no | earlier |
| BBB_Martins | 1,624 | 8, 2000, 0.1, 10 | 431 | no | no | 0.0 | 1 | no | final |
| VDss_Lombardo | 904 | 8, 2000, 0.03, 3 | 431 | no | yes | 0.0 | 1 | yes | final |
| Half_Life_Obach | 532 | 4, 2000, 0.1, 3 | 431 | no | no | 0.8 | 1 | yes | final |
| Clearance_Hepatocyte_AZ | 970 | 4, 1000, 0.1, 3 | 431 | no | no | 0.0 | 1 | yes | final |
| Lipophilicity_AstraZeneca | 3,360 | 4, 2000, 0.1, 3 | 431 | no | no | 0.3 | 1 | no | final |
| hERG | 523 | 6, 500, 0.03, 3 | 200 | yes | no | 0.2 | 1 | no | final |
| CYP3A4_Substrate_CarbonMangels | 535 | 4, 500, 0.1, 10 | 431 | no | no | 0.2 | 1 | no | final |
| CYP2D6_Substrate_CarbonMangels | 532 | 4, 500, 0.1, 10 | 100 | yes | no | 0.0 | 3 | no | final |
| DILI | 379 | 4, 500, 0.03, 1 | 431 | no | no | 0.1 | 1 | no | final |
| Caco2_Wang | 728 | 4, 500, 0.1, 3 | 431 | no | no | 0.0 | 1 | no | final |
| CYP2C9_Substrate_CarbonMangels | 534 | 4, 500, 0.1, 3 | 100 | no | no | 0.0 | 1 | no | final |
| Bioavailability_Ma | 512 | 8, 500, 0.1, 1 | 431 | yes | yes | 0.1 | 1 | no | final |

### 2.7. Independent Chemprop model

As a separate comparison with a message-passing neural network outside the CatBoost selection sequence, Chemprop version 2.3.1^9^ was trained independently on the same five official splits, with a message-passing depth of 3, a feed-forward depth of 2, no dropout, and a maximum of 50 epochs. The hidden dimension was 300, except for `Caco2_Wang`, `AMES`, `LD50_Zhu`, `Solubility_AqSolDB`, and the three CYP inhibition endpoints, for which it was 450, based on hidden dimensions used in earlier runs with hard class labels for the larger endpoints. Inputs comprised the molecular graph, with RDKit two-dimensional descriptors additionally supplied for `hERG`. For each seed, the checkpoint minimizing validation loss was retained; this is the only use of the official validation partition in the present work. Unlike CatBoost, Chemprop used the official validation partition for checkpoint selection; that difference follows from its role as a separate comparator and does not change the official test evaluation.

### 2.8. Leaderboard reporting

Reported leaderboard values use the benchmark aggregation (per-seed metrics rounded to three decimals, then mean and population SD). Table 1 additionally reports the unrounded mean and sample SD. Ranks were computed against a frozen local snapshot of the public leaderboard, rather than by querying it at the time of writing, so that they remain reproducible. Our score for each endpoint was inserted into the snapshot as a new entry and assigned a rank equal to one plus the number of listed scores that are strictly better under that endpoint’s metric; scores that agree to three decimal places therefore share a rank.

`Bioavailability_Ma` was trained and evaluated but not ranked. In the frozen snapshot, the scores listed for this endpoint consist of all sixteen scores listed for `Pgp_Broccatelli` plus four others, and therefore cannot serve as an endpoint-specific reference for ranking. This overlap between the two leaderboard pages has been reported independently as a Therapeutics Data Commons issue.^15^

### 2.9. Training-set size and rank

The association between training-set size and assigned rank was quantified using the Spearman rank correlation coefficient across the 21 rankable endpoints. The difference in training-set size between endpoints ranked first to third and those ranked fourth or lower was assessed using the two-sided Mann–Whitney *U* test. These analyses were specified after the leaderboard results were obtained and are therefore exploratory; the reported *p* values are descriptive and were not corrected for multiple comparisons. Correlation does not imply causation. The endpoints are not statistically independent, because several derive from related assays and overlapping compound sets.

## 3. Results

### 3.1. Held-out performance across the benchmark group

Table 1 reports held-out test performance for all 22 endpoints, and Table 2 the configuration selected for each. Of the 21 rankable endpoints, 7 attained rank 1 on the frozen leaderboard snapshot. Three strictly exceeded the best previously listed score: `Clearance_Microsome_AZ` (Spearman 0.634 versus 0.630), `PPBR_AZ` (MAE 7.349 versus 7.440), and `CYP3A4_Veith` (AUPRC 0.917 versus 0.916). Four matched the best previously listed score to three decimal places and are therefore tied at rank 1: `Solubility_AqSolDB` (MAE 0.741), `HIA_Hou` (ROC-AUC 0.993), `AMES` (ROC-AUC 0.871), and `CYP2C9_Veith` (AUPRC 0.859). A further 5 endpoints attained rank 2 or 3, and 9 attained rank 4 or lower (Figure 2). `Bioavailability_Ma` achieved a ROC-AUC of 0.744 and is reported without a rank (Section 2.8).

**Figure 2.**
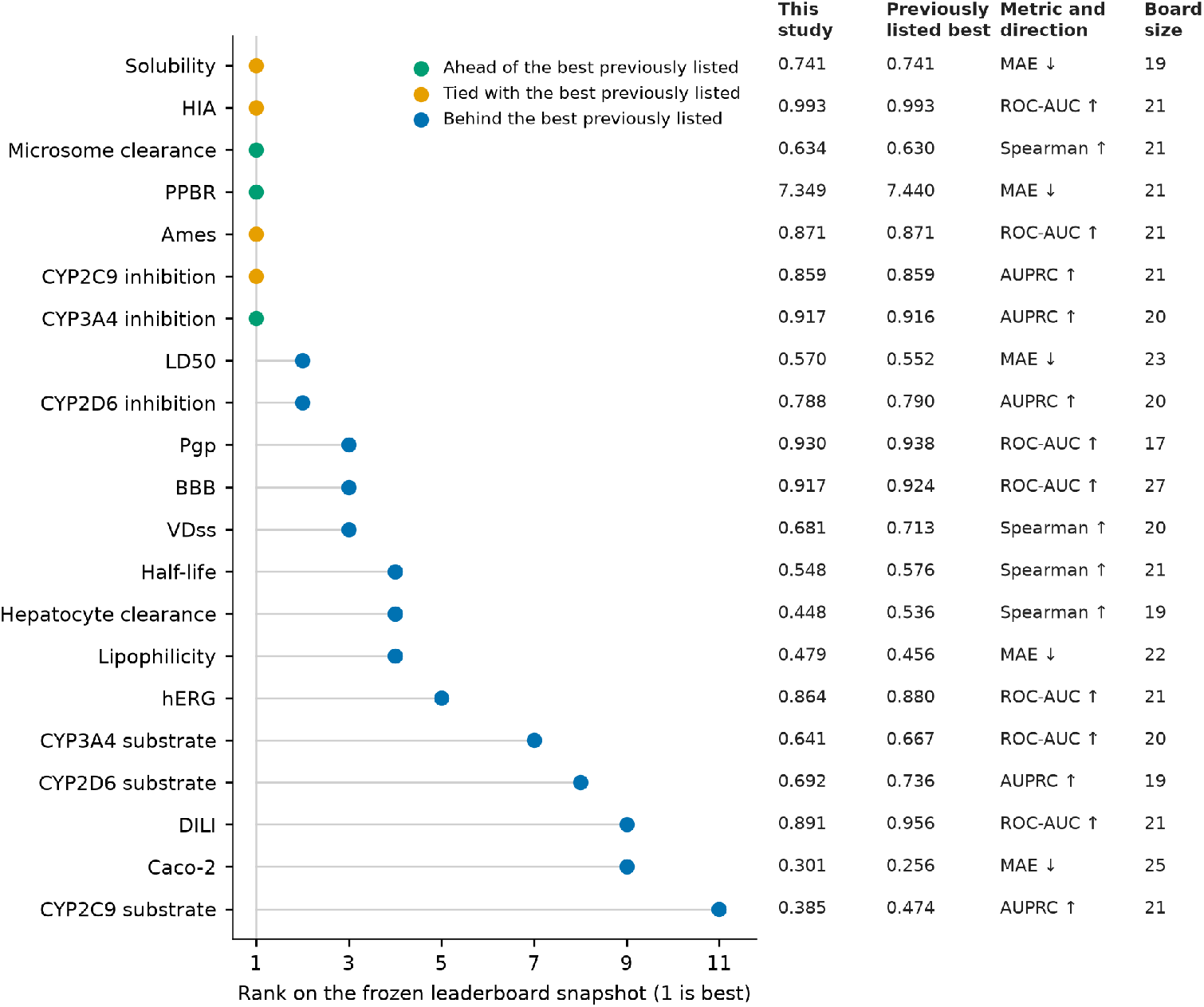
Rank on the frozen leaderboard snapshot for the 21 rankable endpoints, ordered by rank. Green denotes a score strictly exceeding all previously listed entries; orange denotes agreement with the best previously listed score to three decimal places, which also corresponds to rank 1; blue denotes a score inferior to the best previously listed entry. The columns on the right give the leaderboard value, the best previously listed score, the evaluation metric with the direction of improvement, and the number of entries on the board; board sizes differ between endpoints, so ranks are not directly comparable across them. Bioavailability_Ma is excluded (unranked; Section 2.8).

All selected configurations retain the 2,563-dimensional MapLight feature block (Figure 1B); optional components differ across endpoints. The GIN embedding was concatenated at 9 endpoints and omitted at 13 (Figure 3; Table 2). The MiniMol embedding was concatenated to the tree inputs at 5 endpoints (`Bioavailability_Ma`, `Clearance_Microsome_AZ`, `VDss_Lombardo`, `AMES`, and `CYP2D6_Veith`). A non-zero combination weight α was selected at 12 endpoints, with values ranging from 0.1 to 0.8, and α = 0 was retained at 10. Concatenation and α were selected independently. The three-dimensional descriptor block was omitted entirely at 5 endpoints, retained in full at 11, and truncated to between 25 and 200 columns at 6. A positive class weight other than 1 was selected at a single endpoint. In contrast, seven endpoints (`Caco2_Wang`, `BBB_Martins`, `DILI`, `Half_Life_Obach`, `Lipophilicity_AstraZeneca`, `Clearance_Hepatocyte_AZ`, and `CYP3A4_Substrate_CarbonMangels`) retained the initial tree-input feature composition: the MapLight feature block with the full three-dimensional block and neither pretrained embedding.

**Figure 3.**
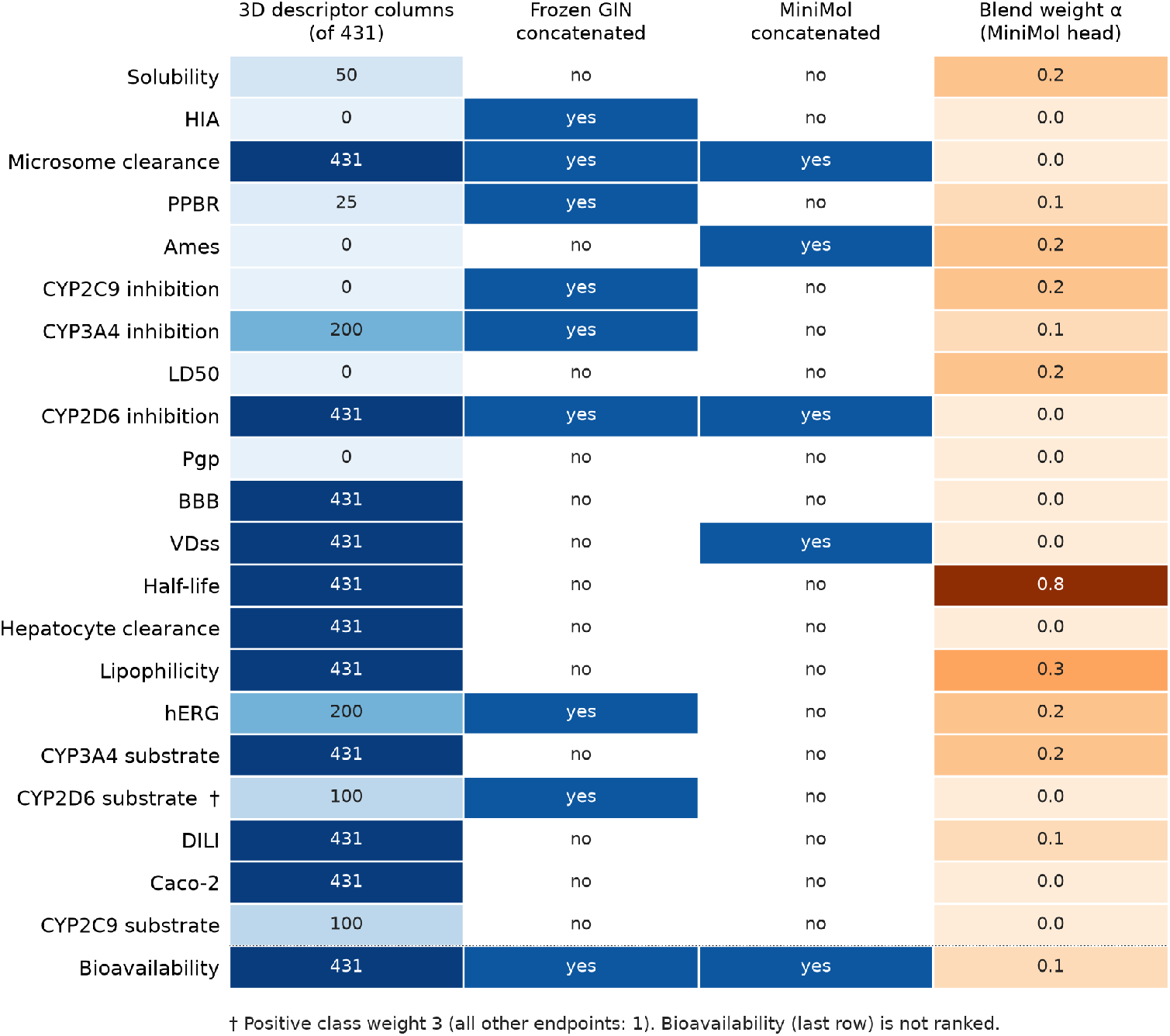
Components selected at each of the 22 endpoints; Bioavailability_Ma (last row) is included but unranked. Columns are shaded independently, without a common scale, and cell annotations give the selected value: the number of three-dimensional descriptor columns retained out of 431, whether the GIN or MiniMol embedding was concatenated to the tree inputs, and the combination weight α of the MiniMol network head. The MapLight feature block, which is identical across endpoints, is not shown. A positive class weight other than 1 (†) was selected only for `CYP2D6_Substrate_CarbonMangels`.

Seven endpoints ranked first: three strictly surpassed the previous best scores, whereas four matched them to three decimal places. All three improvements included the GIN embedding; two also used a non-zero α, and one also concatenated the MiniMol embedding. Among the four matches, two included GIN and two omitted it, while one concatenated MiniMol. Thus, first-ranked configurations differed in their optional selections, with no single optional component consistently associated with rank 1.

### 3.2. Contribution of individual candidate components

The stage introducing the frozen GIN embedding produced the largest changes in held-out performance observed in the selection sequence, concentrated among the CYP inhibition endpoints. At that stage, five-seed test means increased from 0.785 to 0.845 for `CYP2C9_Veith`, from 0.882 to 0.917 for `CYP3A4_Veith`, and from 0.715 to 0.792 for `CYP2D6_Veith`; smaller increases were observed for `hERG` (0.840 to 0.854) and `CYP2D6_Substrate_CarbonMangels` (0.664 to 0.691). These three CYP inhibition endpoints also have the three largest training sets in the benchmark group.

Truncation of the three-dimensional descriptor block produced small changes where it was selected: `HIA_Hou` increased from 0.992 to 0.993 upon complete removal of the block, and `PPBR_AZ` decreased from 7.432 to 7.398 upon retention of 25 columns.

Concatenation of the MiniMol embedding increased `Bioavailability_Ma` from 0.732 to 0.748, `AMES` from 0.862 to 0.870, and `Clearance_Microsome_AZ` from 0.629 to 0.634. Combination with the auxiliary network, applied subsequently with features and hyperparameters held fixed, improved `DILI` from 0.872 to 0.891 (α = 0.1), `Lipophilicity_AstraZeneca` from 0.514 to 0.479 (α = 0.3), and `Solubility_AqSolDB` from 0.765 to 0.741 (α = 0.2). It also improved `Half_Life_Obach` from 0.522 to 0.548 (α = 0.8), `PPBR_AZ` from 7.398 to 7.349 (α = 0.1), and `CYP2C9_Veith` from 0.853 to 0.859 (α = 0.2). α = 1.0 was not selected at any endpoint; for `CYP2C9_Substrate_CarbonMangels`, internal validation AUPRC decreased from 0.329 at α = 0 to 0.154 at α = 1.0.

### 3.3. Internally selected changes that decreased held-out performance

Although some internally selected changes reduced held-out test performance relative to the configurations they replaced, they were retained because they improved the internal validation metric; they are listed below in the order of the selection stages (Section 2.6). For `Bioavailability_Ma`, concatenation of the GIN embedding was adopted despite a decrease from 0.741 to 0.732 and remains part of the final configuration (Table 2). For `hERG`, truncation of the three-dimensional descriptors to 200 columns reduced the score from 0.854 to 0.849, and the 200 columns were retained in the final configuration (Table 2). Concatenation of the MiniMol embedding reduced `VDss_Lombardo` from 0.684 to 0.681 and `CYP2D6_Veith` from 0.792 to 0.788. The combination with the auxiliary network, applied first at α = 0.2, reduced `Bioavailability_Ma` from 0.748 to 0.738 and `hERG` from 0.849 to 0.846. During the subsequent hyperparameter reselection for six endpoints (Section 2.6), the same criterion led to a configuration for `CYP2D6_Substrate_CarbonMangels` that decreased performance from 0.697 to 0.693, whereas `Bioavailability_Ma` rose to the reported 0.744 with a final α = 0.1 and `hERG` rose to 0.864 (combination retained, α = 0.2; Table 2). Those later recoveries do not justify the earlier retentions, which were based only on the internal validation metric at the time.

Finally, adoption of a positive class weight of 3 reduced `CYP2D6_Substrate_CarbonMangels` from 0.693 to 0.692.

### 3.4. Lower ranks and smaller training sets

To place the lower-ranked endpoints in context, we examined how assigned rank relates to training-set size. Nine of the 21 rankable endpoints attained rank 4 or lower (Section 3.1). Rank was inversely associated with training-set size across the 21 rankable endpoints (Spearman ρ = −0.62, p = 0.003; Figure 4; training-set sizes in Table 2). Endpoints ranked first to third had a median of 4,026 training–validation molecules (n = 12), compared with 534 for endpoints ranked fourth or lower (n = 9; Mann–Whitney *U* = 93.0, p = 0.006). Eight of the nine endpoints ranked fourth or lower contain fewer than 1,000 training–validation molecules, the exception being `Lipophilicity_AstraZeneca` (3,360); the smallest is `DILI` (379). Among the seven endpoints at rank 1, six exceed 800 molecules and four exceed 5,000.

**Figure 4.**
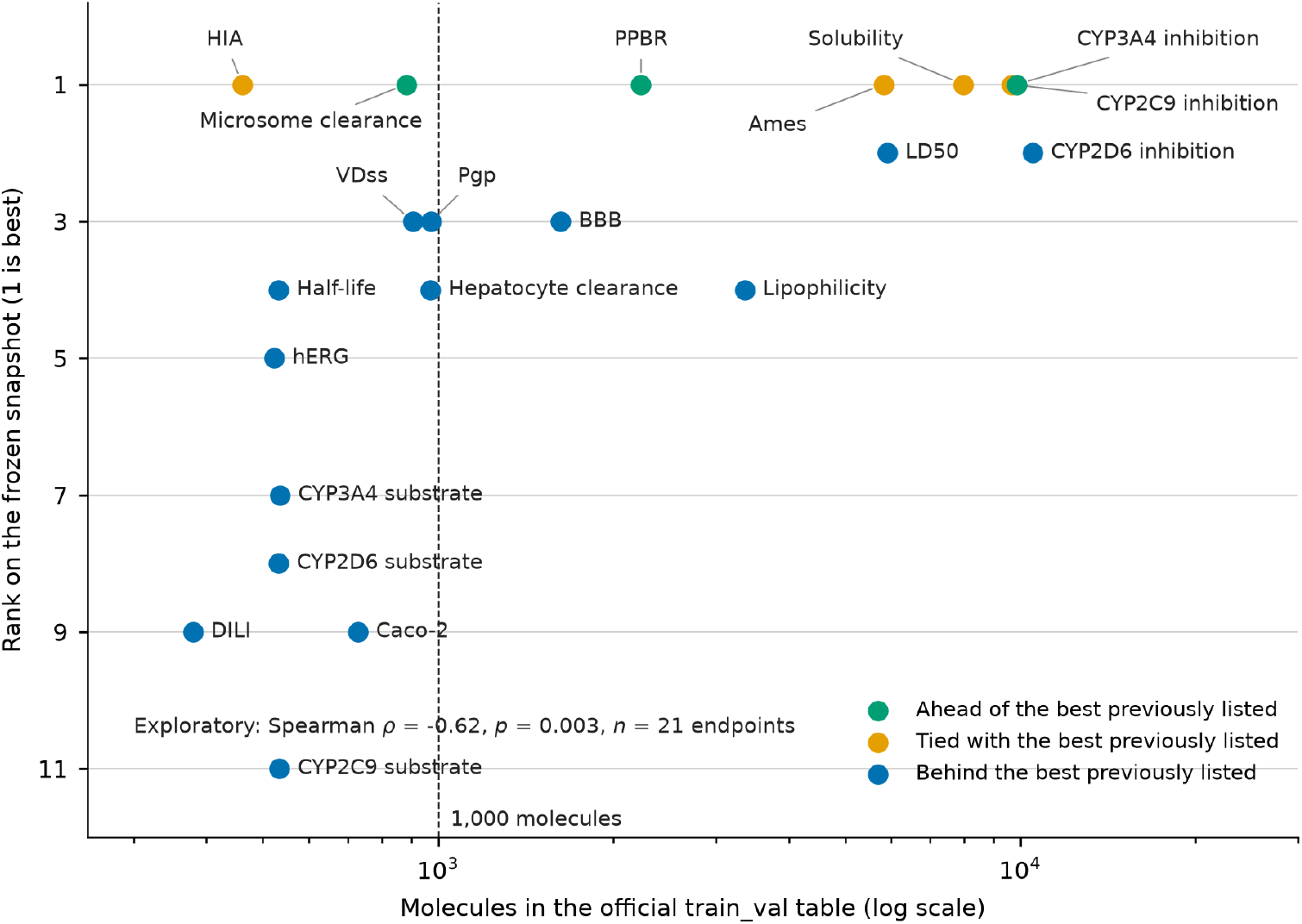
Assigned rank as a function of training-set size for the 21 rankable endpoints. The horizontal axis gives the number of molecules in the training–validation table on a logarithmic scale; the dashed line marks 1,000 molecules. Point colors follow Figure 2. Spearman ρ = −0.62 (p = 0.003); the negative sign reflects that endpoints with more training molecules have numerically smaller (better) ranks. Board sizes differ between endpoints, so ranks are not directly comparable across them. This analysis is exploratory, and training-set size is confounded with the size of the internal validation split used for selection.

The gaps from the best previously listed scores were 0.088 for `Clearance_Hepatocyte_AZ` and 0.089 for `CYP2C9_Substrate_CarbonMangels` in their respective metrics, 0.065 ROC-AUC for `DILI`, 0.045 MAE for `Caco2_Wang`, and 0.044 AUPRC for `CYP2D6_Substrate_CarbonMangels`. These gaps exceed most of the component-wise improvements observed during selection. The lower ranks were associated with smaller training sets and are consistent with, but do not establish, a contribution from limited sample size. As noted in Section 2.9, the size–rank association is exploratory and does not imply causation.

### 3.5. Independent Chemprop model

The main analyses keep CatBoost fixed so that endpoint-specific selection of representations and hyperparameters can be studied under a single learner. This design choice is not a claim that gradient-boosted trees are preferable to message-passing networks on every endpoint. We therefore trained Chemprop on the same five official splits (Section 2.7) and submitted it separately, as a comparator outside the selection sequence. On `Lipophilicity_AstraZeneca`, Chemprop achieved an MAE of 0.412, ranking first among 22 entries in the same snapshot, whereas the selected CatBoost configuration reached 0.479 (tied rank 4; Table 1). The Chemprop predictions were neither used as features for the tree models nor averaged with them. On this endpoint, the message-passing network thus outperformed the selected tree model. This comparison is limited to this endpoint and is independent of the CatBoost results.

## 4. Discussion

Submissions to the TDC ADMET leaderboard commonly apply a single architecture and a single feature set to all 22 endpoints. ^1^ In the present study, endpoint-specific selection under an internal-validation selection protocol yielded a family of models that matched or exceeded the best previously listed score on 7 of 21 rankable endpoints (Section 3.1). Optional components differ across endpoints, while the MapLight feature block is retained throughout.

Fixing any optional choice globally may forgo performance that is attainable at individual endpoints, and making these choices on the held-out test set can bias the estimate of generalization.^8^ The selection protocol was designed to permit endpoint-specific choices while avoiding test-set–based selection. We show that endpoint-specific selection among published components, performed under an internal-validation selection protocol, is a competitive alternative to a uniform recipe on this benchmark. This finding is limited to the benchmark examined here, and whether endpoint-specific selection is preferable more generally remains to be tested.

The components of the candidate set and the choice of a fixed learner were included for specific reasons. The MapLight feature block with CatBoost was adopted as a reproducible baseline that is already competitive under a uniform recipe.^5^ Using it as the fixed learner is a design choice for studying selection under a single learner; it is not a claim that tree ensembles outperform message-passing networks. For this reason, Chemprop^9^ was trained separately and lies outside the selection sequence. On `Lipophilicity_AstraZeneca` it reached an MAE of 0.412 (rank 1), compared with 0.479 (tied rank 4) for the selected CatBoost configuration (Section 3.5). Frozen GIN^14^ and MiniMol^6^ embeddings were included because earlier work indicates that pretrained graph representations can improve performance when kept frozen.^5,6^ Not fine-tuning the encoders keeps the comparison focused on feature selection. Truncated three-dimensional PaDEL descriptors^7^ were included as an optional block that may encode shape- and surface-related information absent from two-dimensional fingerprints. The network trained on the MiniMol embedding alone enters only through a convex combination, so its contribution is an explicit endpoint-wise weight rather than a second, independently tuned model. In the selected configurations (Table 2 and Section 3.1), these components were used differently from endpoint to endpoint, and none of the optional components was selected at all 22 endpoints.

Lower ranks were associated with smaller training sets (Section 3.4), and the endpoints of the TDC benchmark group differ substantially in size.^1^ This analysis is exploratory: the association is correlational, and the endpoints are not statistically independent. Two features of the protocol are relevant. First, with few labeled molecules, improvements from optional components may be difficult to detect; the largest single-stage changes in Section 3.2 occurred at the three CYP inhibition endpoints with the largest training sets. Second, training-set size is confounded with the size of the internal validation split used for selection (approximately 66 molecules for `DILI` and 94 for `CYP3A4_Substrate_CarbonMangels`), so model selection is particularly uncertain for the smallest endpoints. These observations are consistent with a contribution from limited training-set size, rather than the omission of a specific representation, but they do not establish either explanation.

The results should also be read within a limited scope. They should not be read as a reproduction of any single published configuration, since each of those applies one fixed combination of components to every endpoint. In addition, nine endpoints use hyperparameters selected on the initial design matrix rather than on the final one (Table 2), so for these endpoints the hyperparameters were not adapted to the final feature set. Researching these hyperparameters on the final design matrix would address this limitation and is left for future work.

The internal validation split is a single scaffold partition from one seed, and for the smaller endpoints it contains fewer than 100 molecules. On such a split, a small difference between two configurations may reflect noise rather than a true difference in performance. For this reason, the internally selected changes that lowered held-out performance (Section 3.3) reflect adherence to the prespecified rule, not validated improvements. Repeating the scaffold split across several seeds would give a more stable internal criterion, particularly for the smaller endpoints.

Three-dimensional descriptors were computed from a single conformer rather than a conformational ensemble, so properties that depend on conformational distribution are represented only approximately. The pretrained encoders were not fine-tuned, and no predictions or labels from other ADMET models were used. Within the same feature-selection setting, multi-conformer descriptors and limited fine-tuning of the encoders are possible extensions.

## 5. Conclusions

Leaderboard submissions to the TDC ADMET benchmark often apply one architecture and one feature set to all 22 endpoints. This study kept CatBoost fixed and selected configurations separately for each endpoint, using internal scaffold validation. The candidate set comprised published fingerprints and physicochemical descriptors, truncated three-dimensional PaDEL descriptors, frozen GIN and MiniMol embeddings, and a convex combination with a shallow network trained on MiniMol alone. On a frozen leaderboard snapshot, the resulting Ark-ADMET family of 22 models matched or exceeded the best previously listed score on 7 of 21 rankable endpoints and strictly exceeded it on 3. On this benchmark, endpoint-specific selection among published components provided a competitive alternative to uniform recipes without requiring a new architecture. The same approach may offer a practical option for other multi-endpoint benchmarks that apply a single configuration to all tasks.

## Data availability

All benchmark data are publicly available through the TDC ADMET benchmark group. Fitted model weights are available from the Ark-ADMET repository at https://huggingface.co/Eonark/Ark-ADMET.

## Author contributions

Yuzhu Shi: Conceptualization, Methodology, Software, Validation, Formal analysis, Investigation, Data curation, Visualization, Writing – original draft. Haoxi Luo: Conceptualization, Methodology, Validation, Formal analysis, Writing – review & editing. Feng Liu: Resources, Supervision, Project administration, Writing – review & editing. Simiao Zhao: Conceptualization, Supervision, Project administration, Writing – review & editing. All authors reviewed and approved the final manuscript.

